# Multi-tissue DNA methylation age predictor in mouse

**DOI:** 10.1101/119206

**Authors:** Thomas M. Stubbs, Marc Jan Bonder, Anne-Katrien Stark, Felix Krueger, Clock Team BI Ageing, Ferdinand von Meyenn, Oliver Stegle, Wolf Reik

**Affiliations:** Epigenetics Programme, The Babraham Institute, Cambridge CB22 3AT, UK.; European Molecular Biology Laboratory, European Bioinformatics Institute, Wellcome Genome Campus, Hinxton CB10 1SD, UK; Immunology Programme, The Babraham Institute, Cambridge CB22 3AT, UK.; Bioinformatics Group, The Babraham Institute, Cambridge CB22 3AT, UK.; Centre for Trophoblast Research, University of Cambridge, Cambridge CB2 3EG, UK.; Wellcome Trust Sanger Institute, Hinxton CB10 1SA, UK.

**Keywords:** DNA methylation, Epigenetics, Ageing/aging, Ovariectomy, Epigenetic clock, High fat diet, Chronological age, Biological age, Prediction, Model

## Abstract

**Background:** DNA-methylation changes at a discrete set of sites in the human genome are predictive of chronological and biological age. However, it is not known whether these changes are causative or a consequence of an underlying ageing process. It has also not been shown whether this ‘epigenetic clock’ is unique to humans or conserved in the more experimentally tractable mouse.

**Results:** We have generated a comprehensive set of genome-scale base-resolution methylation maps from multiple mouse tissues spanning a wide range of ages. Many CpG sites show significant tissue-independent correlations with age and allowed us to develop a multi-tissue predictor of age in the mouse. Our model, which estimates age based on DNA methylation at 329 unique CpG sites, has a median absolute error of 3.33 weeks, and has similar properties to the recently described human epigenetic clock. Using publicly available datasets, we find that the mouse clock is accurate enough to measure effects on biological age, including in the context of interventions. While females and males show no significant differences in predicted DNA methylation age, ovariectomy results in significant age acceleration in females. Furthermore, we identify significant differences in age-acceleration dependent on the lipid content of the offspring diet.

**Conclusions:** Here we identify and characterize an epigenetic predictor of age in mice, the mouse epigenetic clock. This clock will be instrumental for understanding the biology of ageing and will allow modulation of its ticking rate and resetting the clock in vivo to study the impact on biological age.

## Background

Ageing describes the progressive decline in cellular, tissue and organismal function during life, which ultimately drives age-related diseases and limits lifespan [1]. From a biological perspective, ageing is associated with numerous changes at the cellular and molecular level [2], including epigenetic changes, that is modifications of DNA or chromatin that do not change the primary nucleotide sequence. At present it is not clear which epigenetic changes are causative or correlative, but these mechanisms are of particular interest due to their reversibility, suggesting that rejuvenation might be possible at least in principle [3,4].

Recently, age-correlated DNA methylation changes at discrete sets of CpGs in the human genome have been identified and used to predict age [5-7]. These ‘epigenetic clocks’ can estimate the DNA methylation age in specific tissues [5] or tissue-independently [6] and can predict mortality [8] and time to death [9]. These findings have sparked intense interest regarding the role of DNA methylation in the ageing process and also opened up a number of key questions. Interestingly, while initially designed to predict chronological age, there is evidence that the epigenetic age also reflects biological age and is predictive of functional decline [10-22]. This suggests that the observed methylation signatures might be caused by an intrinsic biological ageing process. One suggestion has been that the methylation clock “measures the cumulative effect of an epigenetic maintenance system” [6], a system that is of critical importance and regulated at multiple levels [23,24]. As such, further insights into the mechanistic properties of this underlying process are of key relevance to understand ageing in more detail and will also be instrumental for the design of future interventions. Consequently, a methylation clock that is applicable to animal models more amenable to experimental interventions would be of considerable importance.

Importantly, while a very small number of age-correlated methylation changes at selected sites in the mouse genome have been reported [25], it is not known whether such an epigenetic ageing clock is conserved between species or a unique property of humans and some closely related primates [6]. Given the general occurrence of the ageing process across the animal kingdom [1], differences in the mechanistic properties of such a clock could explain differences in median lifespan between closely related species. Here we have generated high-resolution methylomes from the experimentally tractable mouse across a wide range of tissues and ages. We find that discrete DNA methylation changes correlate with chronological age and are associated with biological functions. Based on these findings, we generated a multitissue age predictor for the mouse, characterized its properties, and demonstrate that it can be applied to inform other studies by applying it to publicly available datasets, including key biological interventions.

## Results and discussion

### DNA methylation changes in mice correlating with age

In order to study age associated DNA methylation changes in the mouse over a wide range of ages and tissues, we collected liver, lung, heart, and brain (cortex) samples from newborn to 41 week-old mice (Figure 1A). To reduce genetic variability and hormonal variations, we restricted our cohort to male C57BL/6-BABR mice and sampled 3-5 animals per time point. In total we collected 62 samples (Additional File 1) and extracted genomic DNA for methylation analysis from them.

**Figure 1:**
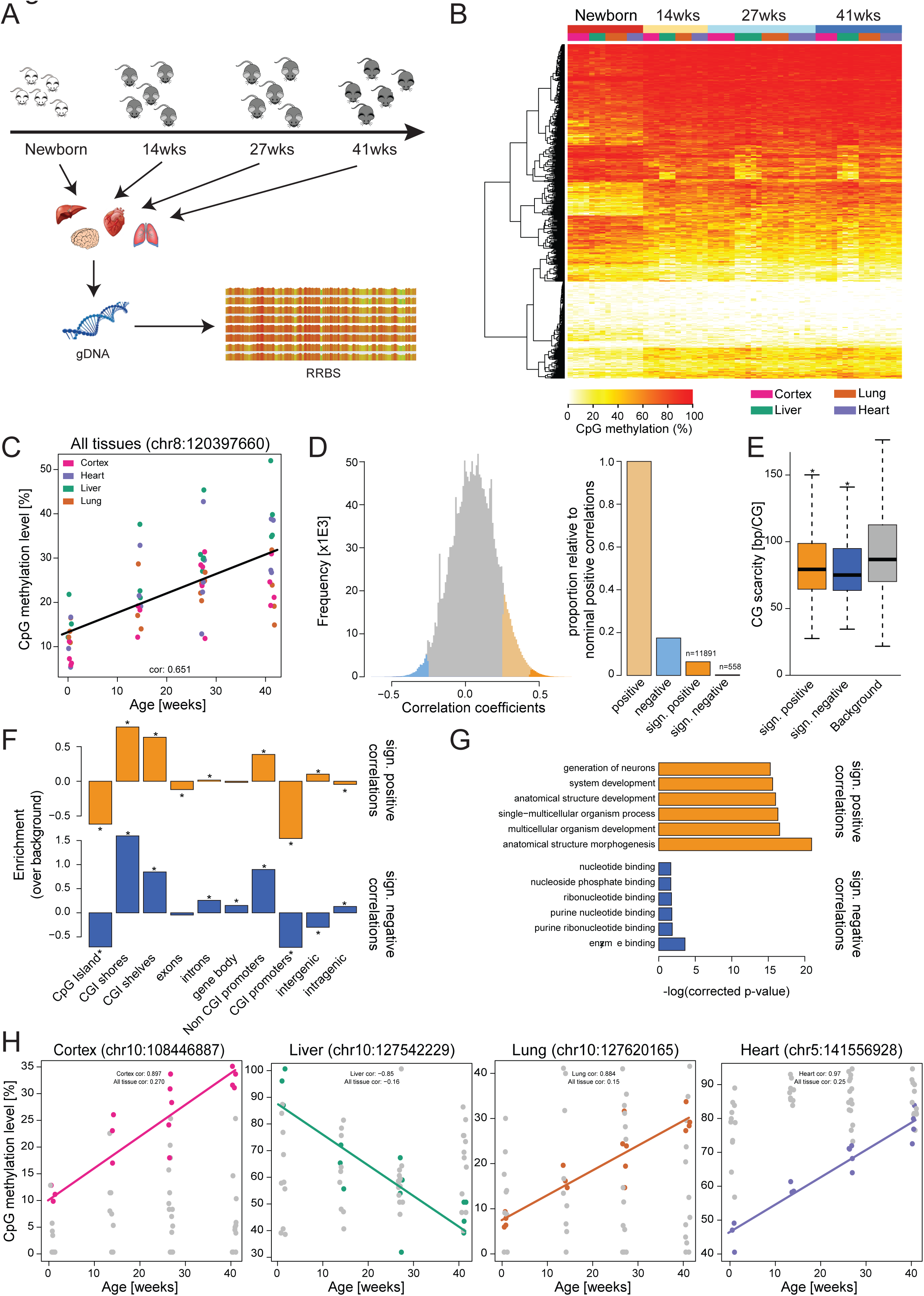
DNA methylation changes correlate with age. (A) Overview of the Babraham dataset. Tissues (liver, lung, heart and cortex) were isolated from mice at 4 distinct time points (newborn, 14 weeks, 27 weeks and 41 weeks). DNA was isolated from these tissues and reduced-representation bisulfite (RRBS) libraries made. (B) Heatmap of the top 500 tissue independent age-associated correlations. Highlighted are ages and tissues, CG sites were clustered by Euclidean distance. (C) Single CG sites within the genome are correlated with age. Shown is an example site (chr8:120397660), with a Spearman correlation with age of 0.65. Tissues are highlighted by colour. Jitter is for aesthetic purposes only. (D) Overview of Spearman correlations calculated over all tissues. The distributions of correlation estimates are shown in the histogram and proportionate numbers of correlations are shown in the barplot. Nominal correlations are highlighted (p-value<0.05) in light orange (positive) and light blue (negative). Significant correlations are highlighted (q-value<0.05) in orange (positive) and blue (negative). Numbers provided above the barplot represent the number of single CG sites within the given category. (E)Surrounding CG content (±500 bp) of the significantly correlated sites. Background was calculated from all CG sites with 5-fold coverage in 90% of samples; CG scarcity is defined as the average CG distance within this 1 kb region. *: Bonferroni corrected p-value<0.05. (F) Enrichment of a correlated CG site falling within a given genomic element. Background was all CG sites with 5-fold coverage in 90% of samples. Tested using binomial test; *: p-value <0.05. (G) GO analysis of the significantly correlated CG sites. GO terms are plotted against −log(corrected p-value). Positive correlations are shown in orange and negative in blue. The 6 most significant GO terms are shown. (H) Tissue-specific Spearman correlations with age. An example is provided of a tissue-specific correlation with age in cortex, liver, lung and heart. Correlations for these CG sites are provided for all tissues combined and for the tissue in question. Jitter is for aesthetic purposes only.

We generated Reduced Representation Bisulphite Sequencing (RRBS) libraries of all samples to be able to assess DNA methylation changes at a wide range of CpG sites and sequenced these to 15x genomic coverage on average. RRBS represents a good compromise between sequencing costs, CpG sites measured and fold genomic coverage obtained. To improve the quantification results, we optimised the standard RRBS library preparation protocol [26] and were able to achieve very low duplication rates and high genomic coverage. On average more than 1.23 million CpG sites in each sample were covered at least 5 fold and of these 0.73 million CpG sites had >5 fold coverage in all samples analysed. Global CpG methylation levels in newborns were around 43% and did not differ significantly between tissues. In the older samples, average methylation levels were slightly higher (~45%) but did not show major differences between ages or tissues (Additional File 2A). Global methylation levels measured by RRBS are generally lower than whole genome bisulfite sequencing estimates, as the method enriches for hypomethylated CpG islands (CGIs) [26]. We also observed low levels of non-CG methylation in all non-brain tissues (Additional File 2B). In agreement with the notion that *de novo* methylation activity in non-dividing cells results in accumulation of CHH methylation, we found that adult cortex samples had higher CHH methylation levels than newborn cortex samples [27]. Together our samples represent the most comprehensive dataset thus far of matched single base resolution methylomes in mice across multiple tissues and ages. Importantly, a hierarchical clustering analysis using Manhattan distances (Additional File 2C) clearly separated the samples by tissue (with the exception of newborn lung samples which clustered together with adult heart samples), highlighting key tissue specific methylation signatures [28].

A correlation analysis showed that DNA methylation at a substantial number of CpG sites across all tissues correlated with age (Spearman’s correlation, with a multiple testing corrected p-value < 0.05) (Figures 1B and C). As expected, the majority of these sites showed age-correlated methylation changes in all or at least 3 tissues (Additional File 2D), suggesting that age-dependent DNA methylation changes at specific sites occur in a coordinated manner across tissues. The correlation values between age and DNA methylation at discrete CpG sites were both positive and negative and normally distributed (Figure 1D). Overall we identified more positive correlations, but this skewing is likely to be the result of the slight global hypomethylation in the newborn samples and represents the general tendency for gain of DNA methylation during development.

To understand whether the underlying sequence composition or genomic context was relevant to the changes observed, we analysed CpG density (Bonferroni corrected two-tailed t-test, p-value<0.05) and genomic context (Binomial test, p-value<0.05) at the significantly correlating sites. While CpGs changing DNA methylation with age were on average in regions with higher CpG density (lower CpG scarcity), the CpG density was not predictive of the methylation changes (AUC = 0.58 or 0.61; see Materials and Methods for details) (Figure 1E). We also found a strong depletion of sites with significant correlations with age at CGIs and CGI rich promoters and conversely a strong increase at CGI-shores and CGI-shelves (4 kb around CGIs) (Figure 1F). This indicates that tightly controlled regulatory regions, such as CGIs, are relatively protected from age associated DNA methylation changes, while regions with intermediate CG density are more prone to changes. A Gene Ontology (GO) analysis of the genes closest (max 4 kb distance) to CpGs with highly significant positive age-dependent changes in DNA methylation revealed a significant enrichment of genes associated with the terms “anatomical structure morphogenesis”, “anatomical structure development” and “developmental process” (Figure 1G). Genes close to sites with significant negative age related correlations were enriched in terms containing nucleotide and enzyme binding. These results suggest that the age-related changes in DNA methylation could alter various important biological processes and future experiments may reveal the regulatory relevance and association with gene expression. In particular, increased DNA methylation at developmentally relevant genes suggests that the ageing process may restrict expression of developmental genes.

We next analysed age-correlations in each tissue independently and identified a large number of sites, which showed exclusively tissue specific changes in DNA methylation with age (Figure 1H and Additional File 2E). Interestingly, only a small fraction of these were shared between 3 or more tissues, indicating that additional age-related DNA methylation changes are characteristic for each tissue and might relate to its intrinsic biological function (Additional File 2F). In particular, GO analysis of genes close to the sites with tissue specific changes revealed highly different GO term enrichments, indicating their unique tissue specific regulation.

The marked differences between the global methylation levels of newborn and adult samples (Additional File 2A) prompted us to also analyse the datasets excluding newborn samples (Additional File 3). A number of sites showed a reversed directionality of age-dependent methylation changes if we excluded the newborn samples from the analysis (highlighted in pink and green; Additional File 3A and B), however the majority of sites (highlighted in blue and orange/red) showed the same correlation characteristics independent of the inclusion or exclusion of newborn samples (Additional File 3C). The correlations between age and DNA methylation in the adult datasets were both positive and negative and the correlation values were normally distributed (Additional File 3D), but none of the correlations were significant (Additional File 3E; Spearman’s correlation, with a multiple testing corrected p-value cut-off of < 0.05). Although the analysis was based on all tissues, only a small number of the age-correlated methylation changes in the adult samples were common to all tissues (p<0.005; Additional File 3F). A subsequent analysis of age-correlations in each tissue independently identified a large number of tissue specific methylation changes (p<0.005; Additional File 3G and H), suggesting that changes in DNA methylation in adults are primarily driven by tissue specific processes, while tissue-independent methylation changes are associated with global developmental processes (Figure 1G).

### DNA methylation levels at a discrete set of CpGs are predictive of age

Having found that methylation at many individual CpG sites did change in an age-dependent manner we decided to generate an epigenetic age predictor in mice (Figure 2A). In addition to our own datasets, we also included previously published datasets comprised of RRBS libraries from liver, lung, muscle, spleen, and cerebellum samples from male and female C57BL/6 mice aged newborn to 31 weeks [29-32]. A description of these datasets can be found in Additional File 4. In short, 129 healthy samples were used to define the training set, with the remaining 189 making up an independent test set, including two datasets that were generated by different labs and hence experimentally independent from the datasets used to train the model. All samples were processed as described in the Materials and Methods. We excluded CpGs located on either of the sex chromosomes or in the mitochondrial genome prior to further processing and analysis.

**Figure 2.**
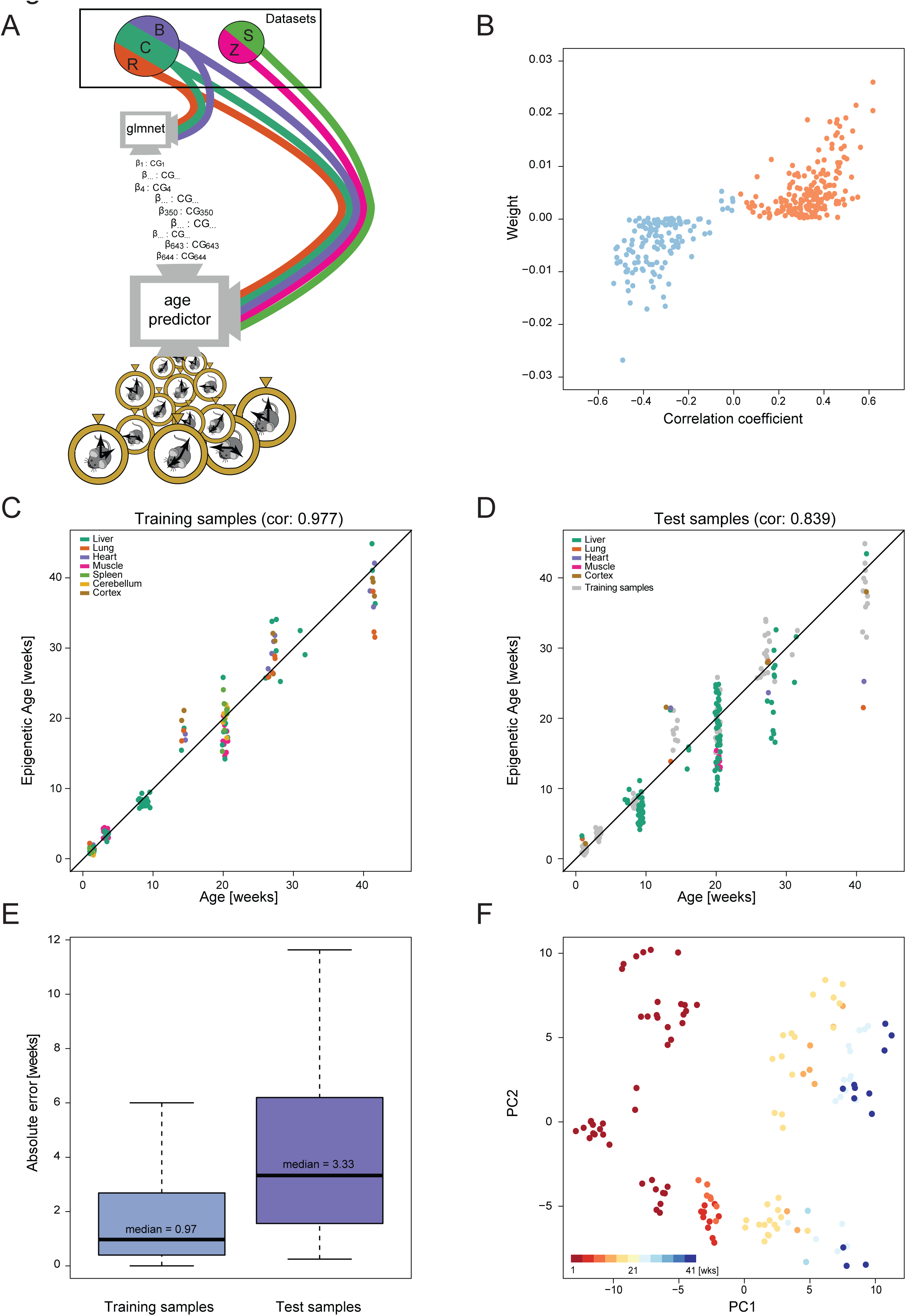
Prediction of chronological age from a mouse epigenetic clock. (A) Flow-chart to illustrate the steps taken in defining the model and testing it. Datasets are displayed as segments of a circle. They are coloured to correspond with later figures, namely: Reizel in brown (R), Cannon in green (C), Babraham in purple (B), Zhang in pink (Z) and Schillebeeckx in light green (S). The two independent datasets are displayed as segments in a separate circle to those datasets utilised for the training phase. The flow of methylation data is shown as colour-coded lines. Training occurs at the node (screen) with the caption: “glmnet”. The chosen CG sites and their corresponding weighting are passed on to the prediction tool itself (node with the caption: “epigenetic age predictor”. Test data enters this prediction tool and age predictions are outputted, as displayed by the pocket watches exiting this node. (B) Scatterplot depicting weight of chosen clock sites against their age-associated correlation, blue are negatively correlated sites and orange are positively correlated sites. (C) Training set ages as predicted by the model, x-axis shows the actual age and y-axis shows the predicted age, coloured by tissue. Jitter is used to represent the experimental error in age estimates. (D) Test set ages as predicted by the model, x-axis shows the actual age and y-axis shows the predicted age, coloured by tissue. (E) Boxplot of the absolute error in the training (left) and test (right) samples. The MAE is indicated. (F) Principal component representation of the sites used in the clock, coloured by age in weeks.

We used an elastic-net regression model to predict log-transformed chronological age, measured in days. Only CpG sites covered by least 5 reads in all samples were used (~18k sites; see Material and Methods for details). Following cross-validation to optimise the model parameters, the final predictor was based on 329 CpG sites (Additional File 5A & Additional File 6); the sites in this predictor will be referred to as mouse (multi-tissue) clock sites. The model selects the most informative sites but allows for some redundancy to increase robustness, and it infers weights for each individual site [33]. The model weights across sites are depicted in Additional File 5B. One implication of this approach is that the clock sites do not necessarily represent the strongest age-correlating sites characterised above (Figure 2B). Similar to the human age predictor described by Hannum et al. [5], the initial starting methylation levels of the mouse clock sites are somewhat predictive of the directionality of their methylation changes with age (Additional File 5C).

As expected, the exponent of the weighted average of the DNA methylation levels of the 329 selected CpG sites was highly correlated with (chronological) age of the individual samples within the training dataset (Figure 2C). Notably, using unobserved test samples, our mouse epigenetic clock was able to accurately predict chronological age in various tissues (Figure 2D) and across multiple independent datasets (Additional File 7A). The accuracy of the model predictions were also independent of sequencing depth of the test samples (Additional File 7B), provided a mean coverage per CpG site of 5 reads or more. This indicates that minor technical variations and coverage differences are well tolerated, and consequently our mouse epigenetic clock model can be applied to a wide range of different settings. This was evident by the fact that the predictor was able to accurately estimate age in two completely independent test datasets [31,32] (Additional File 7C).

The mouse clock performed well across all tissues and ages tested, with an age correlation of 0.839 and median absolute error (MAE) of 3.33 weeks in the test data (Figure 2D and E), corresponding to less than 8.5% error relative to the oldest ages (41 weeks) analysed. In order to compare the accuracy with the human epigenetic clock [6], we calculated the MAE as a proportion of the expected lifespan of a mouse (>100 weeks) and found it to be similar to that reported for the human clock (assuming an average human lifespan of 85 years). Noteworthy, similar to the human clock, the performance of our mouse age-predictor varied between young and old mice. In young animals (<20 weeks) the model-predictions were much more accurate, with a MAE of 2.14 weeks in the test samples. In mice aged 20 weeks or older, the MAE was 4.66 weeks (Additional File 7D). We also attempted to include publicly available whole genome bisulphite sequencing datasets (WGBS), including samples from 24 month old animals [34], to test our age predictor at older ages. Unfortunately, the sequencing depth in most WGBS dataset is significantly lower than in RRBS datasets (mean coverage <5 fold), thus precluding their analysis. It is expected that future high coverage datasets will help to further improve the accuracy of the mouse clock and test its performance at older ages.

To get further insights into the architecture of our multi-tissue age predictor, we performed a principal component analysis (PCA) of the variation within the 329 selected sites in the training datasets. Ninety percent of the observed variability was explained by 69 principal components (PCs) (Additional File 7E), of which 2 PCs (PC1 and PC13) displayed a clear age relation (p<0.05). PC1 captured age-dependent changes and showed a good separation of samples by age; PC2 separated liver samples from the other tissue samples (Figure 2F and Additional File 7F). This analysis highlights that the major variation within the selected CpG sites in the training set is governed by numerous factors, including tissue type and age. However, dataset effects (i.e. technical variations) are not among the major drivers of variation.

Next, we characterized the clock sites in more detail. The CpG sites were distributed across all autosomes with no specific enrichment in any chromosome (Additional File 8A). Similarly to the age-correlated CpG sites, we found a strong depletion over CGIs and CGI promoters but also over non-CGI promoters (Additional File 8B), suggesting that the clock sites were specifically depleted in regulatory regions. CGI shores and intergenic regions showed increased enrichment in clock sites, whereas the CpG density around the clock sites did not show any differences compared to other random CpG sites (Additional File 8C). The 329 clock sites did also not show any specific GO enrichment (not shown), suggesting that the sites selected by the model might not represent a unique biological function and are instead associated with various biological functions.

### Age predictions using the human clock sites in mouse

Given the similarities between our mouse epigenetic age predictor and the previously described human epigenetic clock [6], we asked whether the specific genomic loci described in human could be used to predict age in mouse samples too. First, we attempted to lift-over the genomic locations of the 353 human clock sites to the mouse, by defining regions +/- 500bp around the sites. We were able to lift-over 328 regions, of which 175 regions (in the following referred to as “Horvath clock regions in mouse”) were covered by at least one CpG site in our dataset. Methylation levels at these 175 Horvath clock regions in mouse were only weakly correlated with age (Additional File 9A), which is also true for the Horvath clock sites in human [6].

Next, we assessed whether the 175 Horvath clock regions in mouse could be used for age prediction. Using a ridge model, we were able to generate an age prediction model with a MAE of 11.2 weeks (Figure 3), indicating that the methylation levels at the 175 Horvath clock regions in mouse contain age-related information and are predictive of age in the mouse. The directionality of the mouse-specific and human-specific weighting for any individual CG site within the Horvath clock regions in mouse were completely unrelated (Additional File 9B), suggesting that the human clock [6] is not fully conserved in mouse. Noteworthy, we found that we could also generate age prediction models from matched random regions (see Materials and Methods for details) with an average MAE of 10.6 weeks (Figure 3), highlighting the fact that methylation changes in many genomic regions are age-related and potentially predictive.

**Figure 3.**
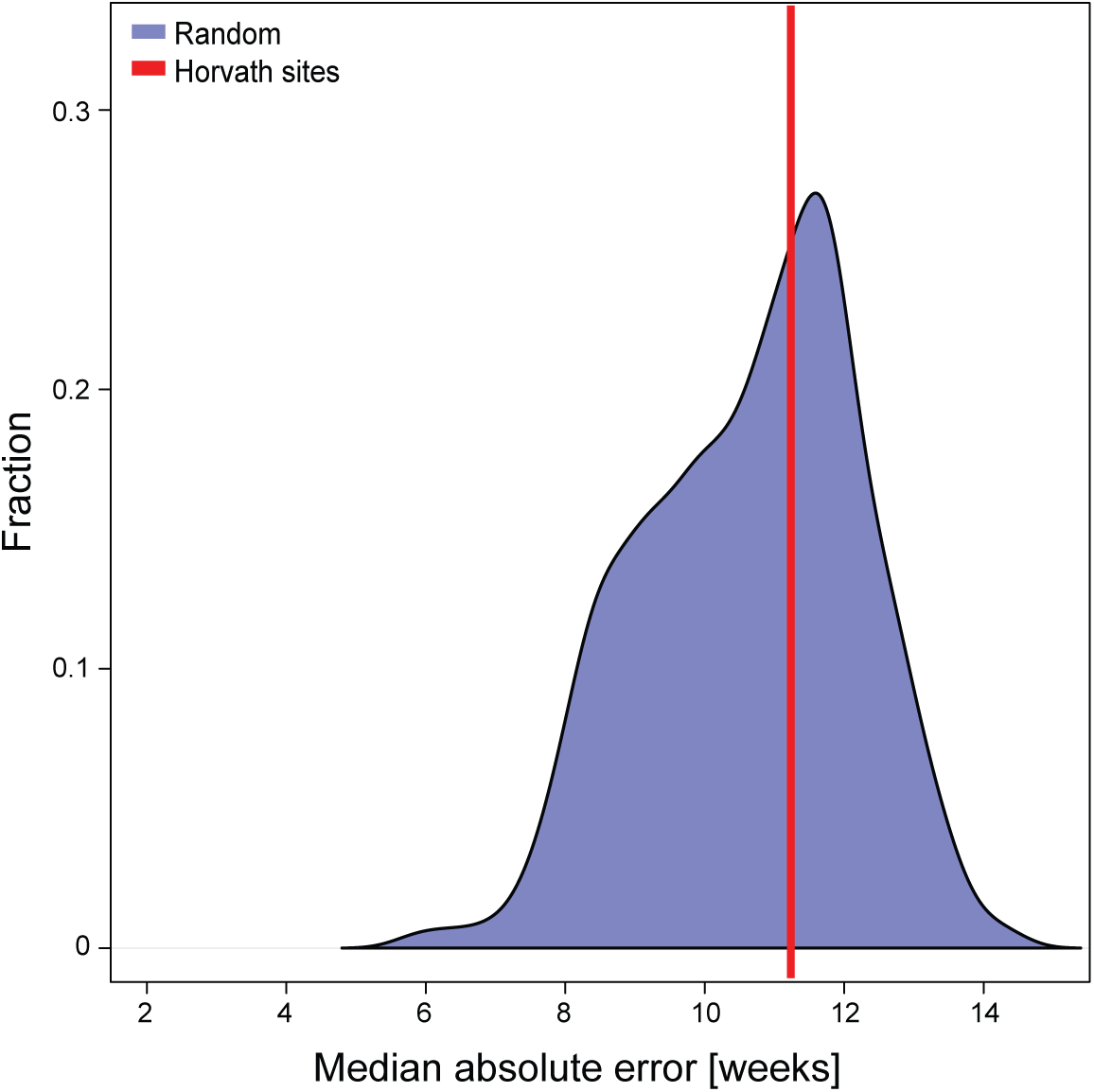
Prediction evaluation of the human Horvath clock sites in mouse. Shown is the MAE of the age prediction model generated using the lifted-over human clock sites [6] (red line). The distribution (blue) shows the MAEs of one thousand age prediction models generated using random selections of 329 regions (see Materials and Methods for further details)

In summary, the predictive accuracy of the sites corresponding to the human clock in mouse samples was significantly lower than our mouse epigenetic clock. These differences are in part due to technical differences in defining the clock sites, but do also highlight species-specific differences, which result in different age-dependent methylation changes at discrete loci and might point to different “ticking” rates of human and mouse ageing.

### DNA methylation age is altered in ovariectomised females and by diet

Given the accuracy of our mouse epigenetic clock to predict chronological age in healthy individuals, we asked whether epigenetic age in the mouse would be affected by gender, diet or other biological interventions. Since it has been reported that DNA methylation shows gender specific patterns [30], we compared the predicted DNA methylation age of female and male mice. Although the training data were predominantly composed of male samples (>72%), we did not find significant differences in the estimated ages between sexes (Figure 4A and Additional File 10), highlighting the robustness of the model and showing that gender specific differences are not skewing the predictions of the mouse epigenetic clock.

**Figure 4.**
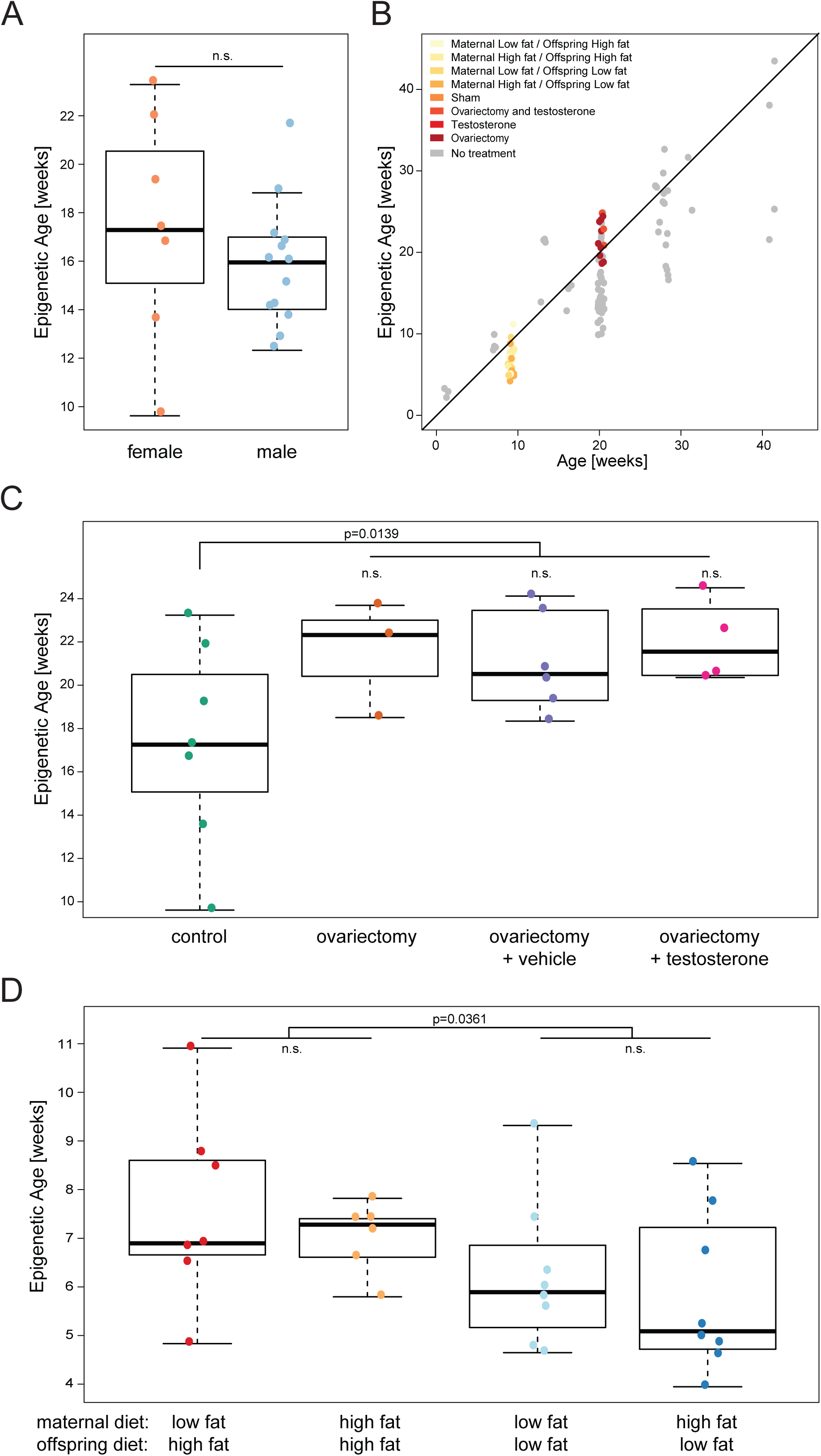
Methylation age is affected by biological interventions. (A) Predicted age of 20 week old liver samples [30], shown separately for males and females. Statistical test performed: t-test, p-value of 0.58. (B) Age prediction in test samples from various biological intervention studies. X-axis shows the actual age and y-axis shows the predicted age, coloured by study. (C) Age prediction in normal female liver samples and samples which underwent an ovariectomy and were administered with either vehicle alone or testosterone [30]. Statistical test performed: Unpaired two-tailed t-test performed to assess the impact of ovariectomy, p-value of 0.014. (D) Age prediction in diet perturbation study [29]. Liver samples from animals with following treatments were analysed: maternal high fat diet followed by adult high fat diet, maternal high fat diet followed by adult low fat diet, maternal low fat diet followed by adult high fat diet and maternal low fat diet followed by adult low fat diet. Statistical test performed: Two-way ANOVA performed, p-values displayed where significant.

We next analysed publicly available RRBS datasets of samples with various biological interventions [29,30]. Overall the epigenetic age predictions were well related to the true chronological age of the individual mice (Figure 4B), but at closer inspection we identified important differences, which depended on treatment.

Reizel et al. performed ovariectomy in females, treated the mice with testosterone (5 mg/mL) or vehicle [30] and analysed their liver DNA methylome by RRBS. Ovariectomy decreases average lifespan in female rats [35], and indeed we found that ovariectomy increased epigenetic age significantly in mice (Figure 4C). Additional testosterone or vehicle treatment had no significant effect on the epigenetic age, and was also shown to not alter lifespan in rats [35]. This suggests that hormonal differences in mice affect ‘biological age’, as measured by our mouse epigenetic clock; similarly in humans breast tissue is found to have accelerated epigenetic ageing [6].

In a 2^nd^ study Cannon et al. [29] characterized the effects of lipid content in maternal and offspring diet on body weight, physiology and DNA methylation status in the liver and reported a significant effect of maternal diet on the susceptibility of the offspring to become obese and develop signs of metabolic disease. The greatest adverse effects were observed when the mothers were fed a low fat diet and the offspring a high fat one. By applying our mouse epigenetic clock to these liver samples, we identified a strong dependency of the predicted epigenetic age on offspring diet (Figure 4D). Offspring on a high fat diet showed accelerated epigenetic ageing, which had a tendency to be further exacerbated if the mothers were fed a low fat diet. This result suggests that biological ageing is modulated by diet and possibly inter- or transgenerational effects and that epigenetic age is a potential powerful measure of biological function.

## Conclusions

In this study, we have generated the thus far most comprehensive set of matched single base resolution methylomes in mice across multiple tissues and ages. This resource allowed us to study the correlation between DNA methylation changes and ageing in detail. Importantly, we were able to establish, for the first time, a mouse epigenetic clock, which estimates age based on the methylation state at 329 discrete CG sites throughout the mouse genome. Our novel epigenetic clock is performing similar to the human epigenetic clock and allowed us to assess (epigenetic) age in unrelated methylation datasets. Noteworthy, the 329 sites of our mouse epigenetic clock perform significantly better in predicting age in mouse samples than the sites in the mouse genome corresponding to the human Horvath clock sites. So far our epigenetic clock has been developed and tested from tissue samples up to 41 weeks of age and future experiments and datasets will be required to assess its accuracy in older mice. The epigenetic clock and the comprehensive set of methylomes are available to the ageing research community and will enable mechanistic and intervention studies in the experimentally tractable mouse model system.

Importantly, we found that the mouse methylation clock is affected by biological interventions, and as such we suggest that the prediction of the clock reflects not only chronological age but also biological age. It will be exciting to test the consequences of manipulations of the ticking rate of the epigenetic clock on biological function, in particular the possible reversibility of ageing associated functional decline. We believe that this work will help scientists gain insights into the biology of ageing and potentially help to discover novel strategies to extend life- and health span in the future.

## Materials and Methods

### Sample collection - Babraham dataset

C57BL/6-BABR male mice were kept under standard conditions in the Babraham Animal Facility. Cortex, heart, liver and lung samples were collected at 4 different ages: newborn (<1 week), 14 weeks, 27 weeks and 41 weeks. All tissues were snap frozen directly after isolation. Genomic DNA was isolated from ~10 mg frozen tissue using the DNeasy Blood & Tissue Kit (Qiagen). A total of 62 samples were collected, processed and further analysed. The resulting dataset are referred here to as Babraham dataset.

### Library preparation

RRBS libraries were prepared from isolated DNA following published protocols [36]. Briefly, RRBS libraries were prepared by MspI digestion of 100 – 500 ng genomic DNA, followed by end-repair and T-tailing using Klenow Exo- (Fermentas). Adapter ligation was performed overnight (homemade adapters) using T4 DNA Ligase (NEB), followed by a cleanup step using AMPure XP beads (Agencourt, 0.9x). Subsequently, libraries were bisulfite treated according to the manufactures instructions (Sigma Imprint Kit; 2 step protocol) and purified using an automated liquid handling robotic system (Agilent Bravo). The libraries were amplified using KAPA HiFi Uracil HotStart DNA Polymerase (KAPA Biosystems), indexing the samples with individual primers. All amplified libraries were purified (AMPure XP beads, 0.8x) and assessed for quality and quantity using High-Sensitivity DNA chips on the Agilent Bioanalyzer. High-throughput sequencing of all libraries was carried out with a 75 bp paired-end protocol on a HiSeq 2000 instrument (Illumina).

### Methylation data processing

For datasets generated in this study, raw paired-end FastQ files were pre-processed to remove the first 13 bp from their 5’ ends, containing unique molecular identifiers (UMI) sequence tags. Both Read 1 and Read 2 UMIs and fixed sequences were written into the read IDs. These trimmed reads were then subjected to adapter and quality trimming with Trim Galore (http://www.bioinformatics.babraham.ac.uk/proiects/trim_galore/; v0.4.2; options: ‐‐paired ‐‐three_prime_clip_R1 15 ‐‐three_prime_clip_R2 15, to also remove potential UMI and fixed tag sequences from the 3’ ends). The trimmed files were then aligned to the mouse genome (GRCm38) using Bismark [37]; v0.16.3, default parameters). The mapped sequences were deduplicated by chromosomal position as well as the UMI sequences of both Read 1 and Read 2 (no mismatches tolerated) using the tool UmiBam (https://github.com/FelixKrueger/Umi-Grinder; v0.0.1; options: ‐‐bam ‐‐ dual_umi). These UMI-deduplicated BAM files were then further processed with the Bismark Methylation Extractor (default parameters) to yield Bismark coverage files.

### Calling of methylation at single CG sites

Mean methylation levels of each CG site covered in each sample were calculated from the Bismark coverage files. In addition, a read count was conducted for each CG site in each sample, so that filtering could be done based on this information in downstream analysis.

### Statistical analysis of age association at single CG sites

For the statistical analysis presented in Figure 1, we first filtered site that had a mean coverage of less than 2 reads or more than 100 reads. This was done to remove spurious reads from library preparation and potential mapping artefacts. For the remaining sites any site that was covered with less than 5 reads in a sample was replaced with NA. Before calculating age association, we further filtered for sites, such that the sites were at least covered in 90% of samples (n = 1,921,569). Ages in days were used when computing the Spearman correlation for each site using the R implementation of the Spearman correlation and multiple testing correction was performed using the Q-value package [38]. For the tissue-specific analysis, a further filtering step was conducted to ensure that there were at least 4 samples being considered for each correlation test. For data exploration we used PCA analysis, on sites (n = 729,785), which were covered in all samples at 5x.

### Predicting significance of age relation based on CpG density

To assess if CpG density is predictive of the methylation age relations we tried to predict if a relation between age and methylation level would be significant based on CpG density. To do so we used CpG density as a linear predictor using multiple thresholds to predict if a methylation age relation would be significant or not using the AUC function for the pROC library [39].

### Genomic enrichment analysis of significant age-associated sites

Normalised likelihood was calculated as:

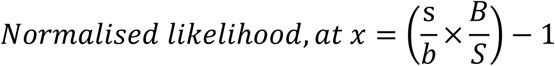
where:

s= number of significant sites at a given x
S = total number of significant sites
b = number of background sites at a given x
B = total number of background sites
CG scarcity
CG scarcity was calculated as:

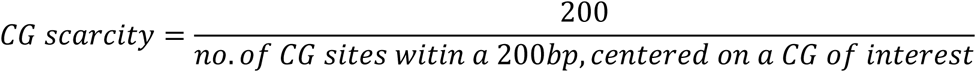

### GO analysis of neighbouring genes

Neighbouring genes were defined for single CG sites that were within 4 kb of a gene. GO terms were defined using the gprofiler online software [40]. For the GO enrichment analysis a background gene list was made consisting of the neighbouring genes (max distance of 4 kb) for all sites considered in that analysis. Significant GO terms were ordered by p-value, and the top 6 GO terms are shown.

### Human-comparative analysis

We defined 1 kb windows around the 21k CpG sites that are interrogated in both the 27k and 450k MethylArray [6], to ensure that the sites could be faithfully lifted-over. These sites were then lifted-over from the human genome to the corresponding regions in the mouse genome (GRCm38). In particular, 91% of the sites selected by Horvath [6] for the human clock were successfully lifted, i.e. 329 of 353. To be able to contrast the human Horvath clock sites [6] to other sites in the mouse genome we chose to use all ~21k sites, of which we were able to lift-over 19k regions. 175 regions corresponding to the 353 clock sites and 10k regions corresponding to all ~21k sites were covered in the Babraham dataset. Adding additional datasets (e.g. Reizel et al. [30]) reduced the number of regions covered dramatically.

Using these sites, we first assessed the age association by comparing the correlation to age in the Horvath clock regions verses random selections of all lifted-over regions. After this we built an age prediction model based on the 175 covered regions corresponding to the Horvath clock sites. For this we built a ridge model as implemented in glmnet, by fixing the alpha parameter to 0. The predictor reaches a MAE of 11.2 weeks. To compare this to background, we built a thousand random models, picking a random set of 329 regions, regardless of coverage, from the 19k regions we could lift over. The average MAE was 10.65 weeks in these random models.

### Predicting age in mice

#### Dataset overview (see Additional File 1)

For defining a more generalizable age predictor, we included four additional external RRBS datasets, which were downloaded from the GeneExpressionOmnibus (GEO) database (https://www.ncbi.nlm.nih.gov/geo/): Reizel et al. [30] (GSE60012; n=173); Cannon et al. [29] (GSE52266; n=40); Zhang et al. [31] (GSE80761; n=4); and Schillebeeckx et al. [32] (GSE45361; n=23). The datasets were processed in the following manner: Raw FastQ files were trimmed with Trim Galore (v0.4.2; parameters: ‐‐rrbs) and then aligned to the mouse genome (GRCm38) with Bismark (v0.16.3; default parameters). The aligned BAM files did not undergo deduplication but were processed directly with the Bismark Methylation Extractor (default parameters) to yield Bismark coverage files.

In the following sections a short description of the dataset is given.

#### Reizel et al. [30]

The Reizel study consists of 173 samples originating from four different tissues: liver, muscle, cerebellum and spleen and data were collected at six time points ranging from 1 to 31 weeks. In the original study gender and tissue specificity of demethylation during ageing has been studied. In the study a perturbation based on castration and restoring testosterone levels after castration has been performed. For the development of the methylation clock the perturbations were not taken into the training, these were left for the test-set. Further information can be found in Additional File 1 and the original publication. After QC there were 143 samples left.

#### Cannon et al. [29]

The Cannon study consists of 40 samples all originating from liver at the age of nine weeks. In the original study the effect of maternal diet on the metabolism of adult offspring was studied. In our study we selected part of the data to be in our training set to reflect the nine-week time point (n=5). The other part of the data is used to assess the effect of diet on ageing. Further information can be found in Additional File 1 and the original publication. After QC there were 36 samples left.

#### Zhang et al. [31]

The Zhang study consists of 4 samples all originating from liver at the age between 6 and 8 weeks. In the original study methylation differences between different strains of mice, and the difference between mouse and zebrafish DNA-methylation levels are assessed. In our study these samples were used as a validation to see how the predictor works for an unobserved time-point. The age of these mice has been set to 7 weeks in our study. Further information can be found in Additional File 1 and the original publication. After QC there were 4 samples left.

#### Schillebeeckx [32]

The Schillebeeckx study consists of 23 samples all originating from the liver, the adrenal gland and from endometrial cancer. The mice were ovariectomized at the age of 3 to 4 weeks, samples were collected after an additional 3 months. In the original study a laser capture microdissection RRBS method was introduced. For our study we selected the liver samples, which were generated using normal RRBS, and after QC 3 samples were left. In our study these samples were used as a validation to see how the predictor works for an unobserved time-point. The age of these mice has been set to 16 weeks in our study. Further information can be found in Additional File 1 and the original publication.

#### Age prediction

To predict the mouse age, we adopted a similar approach to those utilised in human studies [5,6], namely an elastic-net regression model. Firstly, we intersected the sites, which were available with more than 5-fold coverage in all training and test samples used, totalling to 17,992 sites. By selection of the sites available in all datasets we hope to have selected a set of methylation sites, which will be present in most RRBS studies, irrespective of size selection and data handling. In addition, we filtered out both sex chromosomes (X and Y) and the mitochondrial genome to ensure that the model would not be sex-specific nor hampered by the unreliability of mitochondrial genome bisulfite conversion. After selection of the sites and samples, we have used a quantile normalization to normalize the methylation values, followed by a standardization, putting the mean methylation per site to 0 and the standard deviation to 1.

For the predictor we used the elastic-net generalized linear model as implemented in the GLMNET package [33]. To optimize the alpha, defining the elastic net mixing parameter (1 for lasso to 0 for ridge) and to optimize the lambda, the regularization parameter, we used a double-loop cross-validation set-up. This setup is described in Ronde et al. [41]. We have trained the model to predict the log transformed mouse age (in weeks); 3 weeks were added before the log transforming the ages, to be able to predict samples pre-birth.

For the training set we selected 129 healthy samples from the Babraham, Reizel and Cannon study, sample details are in Additional File 4. By using an internal 10 fold cross-validation, in the inner loop the optimal alpha (0.05) and optimal lambda (0.93) were identified. In the outer loop the actual performance of the predictor was scored (as assessed by the mean squared error). After the cross-validation in the training we built the final model on all 129 samples to get our final model. To get to our final model we have taken the beta’s as derived from GLMNET for the selected sites (329) and trained a quadratic function using the nls function in R to transform the raw prediction scores (sum of the product of the beta weights multiplied by their respective methylation level) to the log age in weeks. The final function used is:

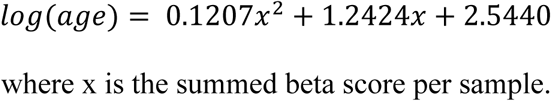

A set of healthy and treated samples, originating from the same three studies, and the Schillebeeckx and Zhang study were used to assess the usability of the final model. The MAE of the prediction was found to be 3.33 weeks. Furthermore, the model has been used to assess the influence of diet on the methylation age, using the Cannon training samples and the influence of male and female castration on the methylation age.

MouseEpigeneticClock GitHub project:

To predict methylation age from new samples we have generated an easy to use R project and deposited it as a GitHub project:

https://github.com/EpigenomeClock/MouseEpigeneticClock

## Abbreviations

RRBS: Reduced Representation Bisulphite Sequencing
WGBS: Whole Genome Bisulphite Sequencing
GO: Gene Ontology
PCA: Principal Component Analysis
MAE: Median Absolute Error
CGI: CpG Island

## Declarations

### Ethics approval

All animal work carried out as part of this study is covered by a project license (to W.R.) under the 1986 Animal (Scientific Procedures) Act, and is further regulated by the Babraham Institute Animal Welfare, Experimentation, and Ethics Committee.

### Accession Numbers

The accession numbers for the next-generation-sequencing data reported in this study are GEO: GSE93957.

### Authors’ contributions

TMS, FvM and WR designed the study; TMS, FvM and AK-S collected samples and performed experiments; TMS, MJB and OS analysed data and developed the model; FK performed processing of data; TMS, MJB, FvM, OS and WR interpreted data and wrote the manuscript. All authors approved the final manuscript.

## Acknowledgements

We thank all the members of the Reik and Stegle laboratories for helpful discussions. In particular we thank Poppy Gould and Wendy Dean for scientific advice and support, Kristina Tabbada for assistance with high-throughput sequencing, and Anne Segonds-Pichon for statistics advice.

## BI Ageing Clock Team (alphabetical)

Daniel Bolland^a^
Geoff Butcher^b^
Tamir Chandra^c,f^
Stephen J. Clark^c^
Anne Corcoran^a^
Melanie Eckersley-Maslin^c^
Lucy Field^c^
Ulrika C. Frising^b^
Joana Guedes^b^
Irene Hernando-Herraez^c^
Jon Houseley^c^
Fiona Kemp^d^
Amy MacQueen^e^
Klaus Okkenhaug^e^
Martyn Rhoades^d^
Milou J. C. Santbergen^b^
Marisa Stebegg^b^
Marc Veldhoen^b,g^

## Footnotes

a Nuclear DynamicsProgramme, The Babraham Institute, Cambridge CB22 3AT, UK

b Lymphocyte Signalling Programme, The Babraham Institute, Cambridge CB22 3AT, UK

c Epigenetics Programme, The Babraham Institute, Cambridge CB22 3AT, UK

d BSU, The Babraham Institute, Cambridge CB22 3AT, UK

e Immunology Programme, The Babraham Institute, Cambridge CB22 3AT, UK

f Current address: MRC Institute of Genetics & Molecular Medicine, The University of Edinburgh, Edinburgh EH4 2XU, UK

g Current address: Instituto de Medicina Molecular, 1649-028 Lisboa, Portugal

## Additional material

**Additional File 1:** List of all samples collected in this study, as well as all additional publicly available datasets used. (Excel Format; AdditionalFile1.xls)

**Additional File 2: Characterisation of ageing associated methylation changes at discrete CpGs**, related to Figure 1. (PDF Format; AdditionalFile2.pdf)

**Additional File 3: Characterisation of ageing associated methylation changes at discrete CpGs excluding newborn samples**, related to Figure 1. (PDF Format; AdditionalFile3.pdf)

**Additional File 4: Table of Training and Test samples**. (Excel Format; AdditionalFile4.xls)

**Additional File 5: Properties of the mouse epigenetic clock sites**, related to Figure 2. (PDF Format; AdditionalFile5.pdf)

**Additional File 6: Table of mouse epigenetic clock sites**. (Excel Format; AdditionalFile6.xls)

**Additional File 7: Additional detail on the mouse epigenetic clock**, related to Figure 2. (PDF Format; AdditionalFile7.pdf)

**Additional File 8: Information on the datasets properties of the clock sites**, related to Figure 2. (PDF Format; AdditionalFile8.pdf)

**Additional File 9: Testing age relation of the human clock sites in the mouse**, related to Figure 3. (PDF Format; AdditionalFile9.pdf)

**Additional File 10: Predicted age of test samples as coloured by sex**, related to Figure 4. (PDF Format; AdditionalFile10.pdf)

